# Proprioceptive Cortical Neurons Implement Optimal State Estimation

**DOI:** 10.64898/2026.02.23.707432

**Authors:** Melanie Palacio-Manzano, Irina Scheer, Mario Prsa

## Abstract

While the primary somatosensory cortex (S1) is known to be essential for skilled limb movements, it remains unclear how distinct functional ensembles within its network uniquely contribute to motor control. By isolating a stable population of proprioceptive S1 neurons in layer 2/3, we demonstrate how their selective removal impacts goal-directed reaching. We found that following their microablation global reach kinematics remained stable, yet natural movement variability was reorganized. Trajectories became more spatially dispersed while simultaneously becoming more geometrically stereotyped. Through computational modeling, we show that this seemingly paradoxical increase in one form of variability alongside a reduction in another reflects a failure in optimal state estimation. In contrast, broad S1 lesions mask the unique contribution of this ensemble, revealing instead that cortical feedback is temporally phased, engaging during high-precision grasping and retrieval rather than initial reaching. These findings identify a functionally defined subset of cortical neurons as the biological substrate for a core postulate of optimal control theory, operating within a broader S1 circuit specialized for fine motor refinement.

## Introduction

The primary somatosensory cortex (S1) encodes proprioceptive feedback signals^1–6^ that seem to be essential for executing skilled, goal-directed limb movements. Causal investigations into S1’s role in motor control have relied on broad regional perturbations, such as focal lesions^7–11^, pharmacological inactivation^12^ or optogenetic silencing^13^, and demonstrated its critical contribution to behavior^14^. However, S1 encodes heterogenous modalities^1,15,16^, including proprioceptive, tactile, and thermal signals, each across functionally distinct subnetworks of neurons. For example, layer 5 corticospinal neurons^8^ and layer 2/3 cortico-cortical projection neurons^17^ likely play divergent roles in sensorimotor integration and control. Global cortical perturbations affecting the entire network therefore inevitably mask the unique contributions of specific functional groups. Determining these unique contributions will reveal the computational logic of local cortical circuits and how they process proprioceptive feedback^18^.

Targeted single-cell ablation or holographic activation experiments have demonstrated that a small number of functionally defined neurons in sensory cortices can shift sensory representations and perception^19–25^. These findings clarify how functional neuronal ensembles mediate perceptual aspects of behavior. In contrast, it remains unclear how analogous ensembles in S1, particularly proprioceptive signals within layer 2/3 neurons, regulate motor execution. It is unknown if and how the loss of these functional units would manifest in the kinematics of a complex movement. Do they primarily convey perceptual information about the body in space^1^, or do they also provide stable signals for feedback control of its movements?

To address these questions, we imaged the activity of layer 2/3 neurons in the mouse forelimb somatosensory cortex (fS1) and identified a subset with stable proprioceptive responses to passive limb displacements. Selective photoablation of a small number of these proprioceptive neurons produced an unexpected effect on limb trajectories during execution of reach-to-grasp movements. Movements started following a more stereotyped trajectory across trials but at more dispersed spatial locations, while overall kinematics remained relatively stable. Remarkably, an Optimal Feedback Control (OFC) model^26^ replicated these experimental findings by simulating a suboptimal integration of external sensory feedback with internal prediction, suggesting that this specific neuronal ensemble acts as an optimal state estimator. We next demonstrate that broad fS1 lesions mask this unique contribution by inducing widespread kinematic and spatial distortions. In both cases, classifying deficits across sequential movement components reveals a temporally phased engagement of fS1 in feedback control. Together, our results provide experimental evidence for a core postulate of the OFC theory, which is that noise dependent feedback drives the variability of unique movement trajectories that in its absence default to a stereotyped pre-planned path^27,28^. We identify layer 2/3 proprioceptive neurons as key functional units, within a broader fS1 circuit, that enable such behavioral flexibility through optimal state estimation.

## Results

### Stable cortical proprioceptive tuning

To provide consistent feedback for motor commands, proprioceptive neurons should maintain a stable functional identity. However, representational drift, the spontaneous or experience-dependent shifting of neuronal tuning, is a hallmark of many sensory and higher-order brain regions^25,29–31^. Can individual proprioceptive neurons in somatosensory cortex provide a consistent feedback signal over time, or must stability instead be achieved at the population level through compensation for tuning drifts^32,33^? To answer this question, we first examined the longitudinal stability of proprioceptive neurons to determine whether they preserve their functional properties throughout repeated sensorimotor experience.

Following our established protocol^1^, we delivered proprioceptive stimuli via passive displacements of the mouse forelimb (Fig. 1A). Using two-photon calcium imaging, we longitudinally tracked the responses of identified neurons in the forelimb somatosensory cortex (fS1) of transgenic mice stably expressing GCaMP6f in L2/3 neurons, across intervals ranging from 9 to 72 days (Fig. 1B). Between these imaging sessions, mice underwent continuous sensorimotor training consisting of both the passive displacement task (264 ± 63 trials/session, mean ± s.d.) and an active water droplet reaching task^34^ (404 ± 273 reaches/session, mean ± s.d.). These training modalities were alternated throughout the inter-imaging periods, occurring on average on 43% and 21% of the intervening days for passive and active tasks, respectively (8 mice, 30 sessions). Consequently, the duration of the inter-imaging intervals was linearly correlated with the total amount of sensorimotor experience, allowing us to evaluate its impact on the tuning stability of the tracked fS1 neurons (Fig. 1B).

**Figure 1.**
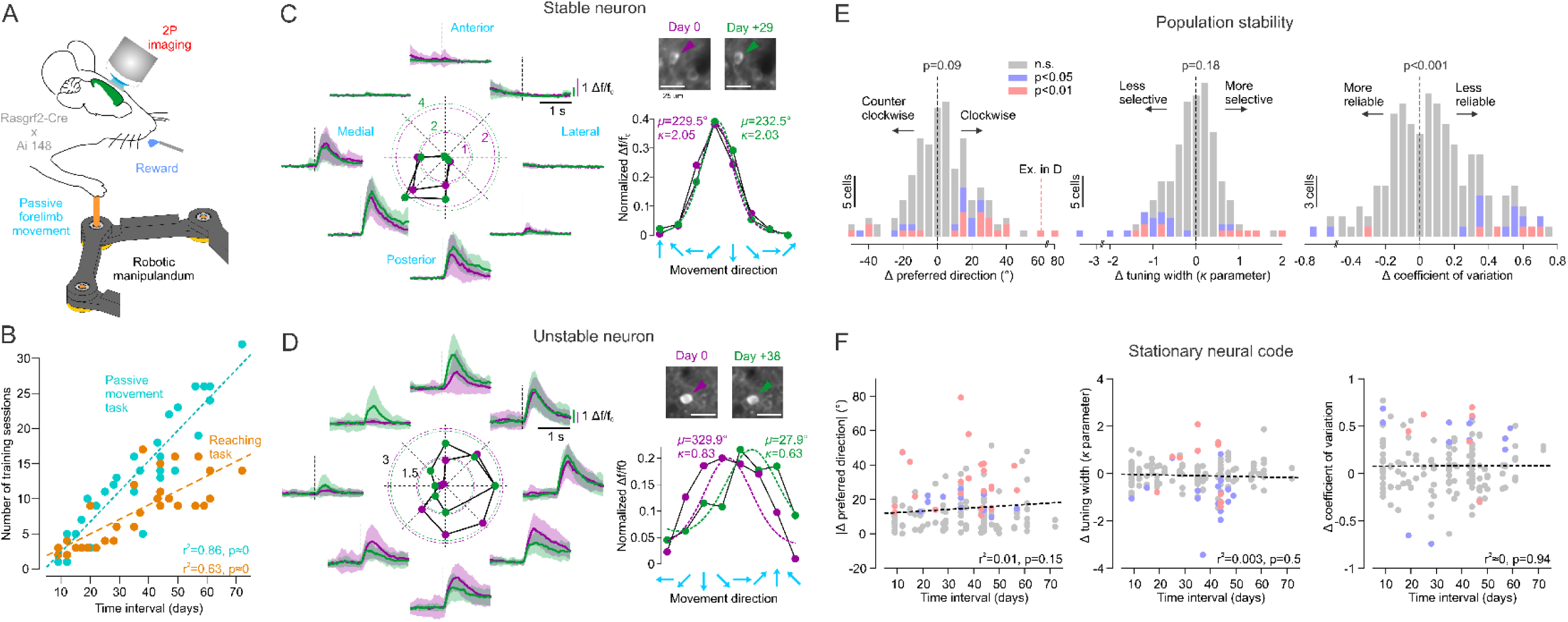
Stable representation of proprioception by fS1 neurons. **A**: Schematic of the passive limb displacement task. **B**: The number of passive and active training sessions scaled linearly with the temporal interval between imaging sessions used for tuning stability analysis (N=30 sessions, 8 mice). **C**: Example of a proprioceptive neuron with stable directional tuning. Left: mean (±s.d.) responses to 8 directions of passive limb displacement on the first imaging day (magenta) and 29 days later (green). Polar plot: peak responses as a function of movement direction (data are peak ΔF/F_0_ post-stimulus activity). Right, bottom: Von Mises fits (dotted lines) to the normalized peak responses tuned to movement direction on the two imaging days. μ and κ parameters of the Von Mises fit quantify the preferred direction and tuning selectivity, respectively. Right, top: cropped session average two-photon images showing preserved identity of the tracked neuron (arrows). Scale bars: 25 μm. **D**: Same data as in C for an example neuron with unstable directional tuning. **E**: Distributions of changes in preferred direction (μ parameter), tuning selectivity (κ parameter) and coefficient of variation between the early and late imaging days of the tracked neurons (N=85, 8 mice). Individual neurons with significant changes (bootstrap test) are highlighted in red (p<0.01) and blue (p<0.05). Population level preferred direction (p=0.09. t-test) and selectivity (p=0.18, signed rank test) remained stable, while the coefficient of variation increased (p<0.001, t-test). **F**: Changes in response tuning or reliability did not correlate with the amount of sensorimotor training (same color code as in E).

Across 8 mice and 30 imaging fields of view, we first assessed the longitudinal stability of the proprioceptive population. Of the 116 neurons identified during initial imaging, 92 remained responsive in the second session (≈20% functional dropout), while 5 of the 97 neurons identified in the second session were newly recruited (≈5% drop-in).

To characterize directional tuning dynamics within the stable population (N=92), we tested eight coplanar directions of forelimb displacement. The neurons’ preferred directions and directional selectivity were quantified using the μ and κ parameters, respectively, derived from a Von Mises fit to the peak average ΔF/F_0_ responses. We subsequently restricted our stability analysis to neurons with significant Von Mises fits (N=85), classifying them as either stable (Fig. 1C) or unstable (Fig. 1D) based on significant shifts in their tuning parameters (p<0.01, randomization test). Because each neuron was tested across two displacement sets (centrifugal and centripetal, see Methods), each unit contributed two independent data points to the overall analysis. Instability in preferred direction and directional selectivity was observed in 23.7% and 17.3% of units, respectively (Fig. 1E). Notably, while population-level selectivity remained unchanged over time, the reliability of these evoked responses diminished, as evidenced by a significant increase in the coefficient of variation (Fig. 1E). Finally, we asked if sensorimotor experience drove these changes in tuning and reliability. In all cases, the magnitude of the fluctuations did not correlate with the time interval between imaging sessions (Fig. 1F), suggesting that they reflect intrinsic circuit turnover rather than experience-dependent plasticity.

We conclude that fS1 proprioceptive neurons largely retain stable functional properties. Changes within the unstable minority were typically subtle; the example in Fig. 1D represents the second largest shift in our dataset (Fig. 1E), as most significant tuning changes did not exceed 40°. The stability suggests that these neurons participate in a fixed sensory reference frame, potentially providing a consistent feedback signal for motor control.

### Proprioceptive fS1 neurons support the natural variability of reaching paths

If proprioceptive fS1 neurons indeed serve as a fixed feedback source for the motor system, the execution of active movements should depend on the integrity of this specific ensemble. In a new set of experiments, we therefore selectively removed these neurons using targeted microablation^21,35,36^ to determine how the loss of a stable sensory signal modifies reaching performance.

We first identified proprioceptive neurons activated by passive limb displacement, before transitioning to the active reaching task. In this established behavioral framework^34^, freely moving mice were trained to reach for water droplets positioned at a distance that required forelimb extension but precluded licking. This motor skill is performed almost spontaneously and provides a controlled context for assessing motor execution. Once mice achieved baseline performance, we tracked 3D paw trajectories using a stereo camera system (Video 1, Fig. 2A, see Methods), both before and after ablating the identified proprioceptive neurons in the contralateral cortex. Photoablation was executed by focusing and scanning the imaging laser over individual neuron soma at high power. Successful ablation was confirmed by an abrupt and irreversible increase of nuclear calcium fluorescence, an established marker of apoptosis^37,38^, while adjacent neurons remained unaffected. Functional loss was verified by subsequent absence of both spontaneous and stimulus-evoked calcium transients (Fig. 2B). Such selective ablation of a small subset of functionally defined layer 2/3 neurons attenuates responses across the broader population of spared neurons, specifically those sharing the same stimulus representation^21^. Conversely, ablating silent unrelated neurons does not appear to alter the encoding properties of functional ensembles^21^.

**Figure 2.**
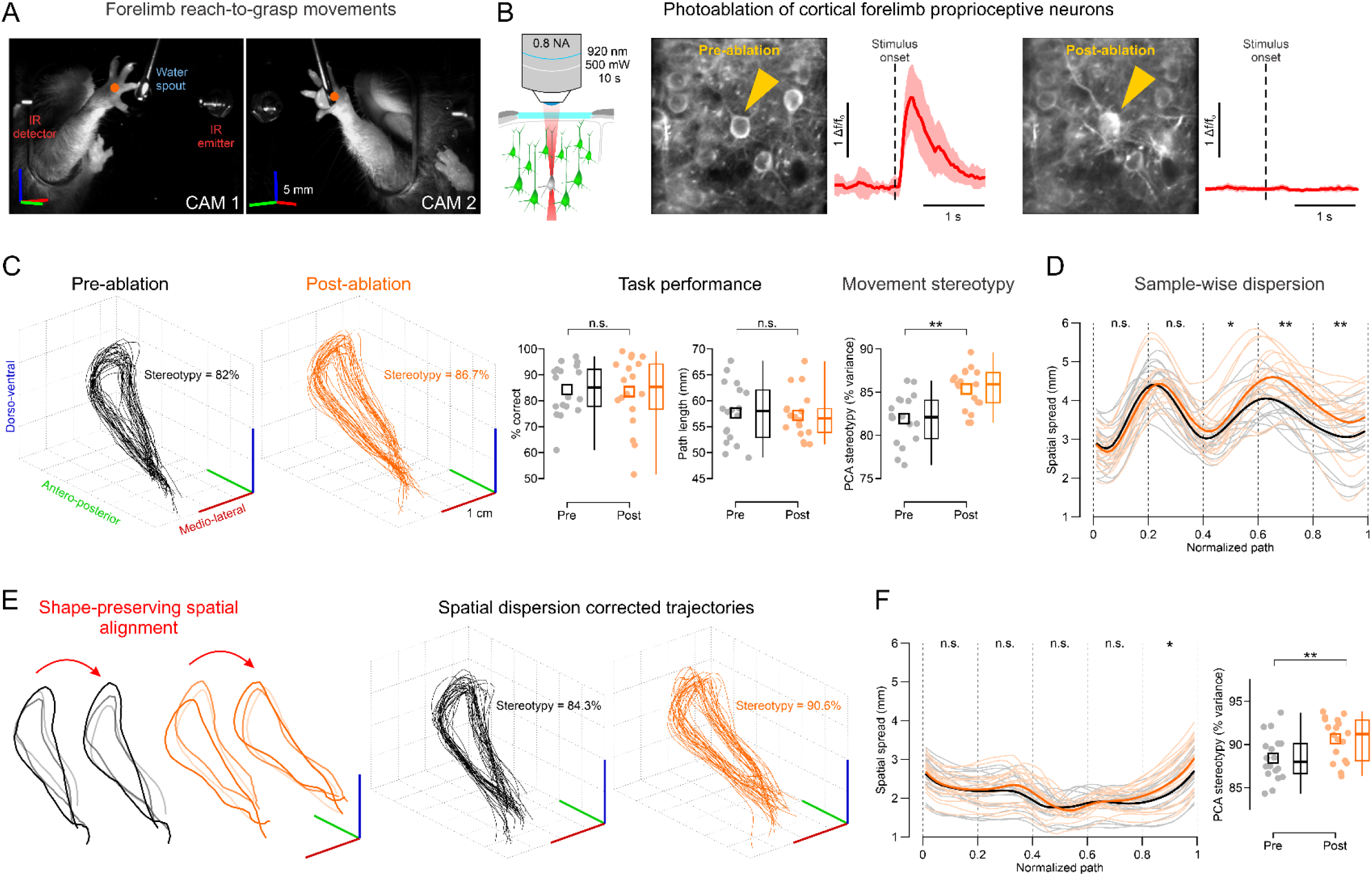
Changes in reaching trajectories after selective microablation of proprioceptive fS1 neurons. **A**: Still frames of the stereo camera system tracking the mouse paw during the reach-to-grasp task. **B**: Schematic of single cell photoablation (left) and example two-photon images (maximum projections) depicting a proprioceptive neuron (arrows) pre- and post-ablation (right). Red traces are corresponding mean (±s.d.) responses to passive limb displacement in the neuron’s preferred direction. **C**: Pre- and post-ablation 3D movement trajectories of example sessions from the same mouse (left, only the first 25 trials are shown for clarity) with stereotypy index values. Right: comparison of correct performance, trajectory path length and stereotypy pre- and post-ablation (N=18 sessions, 6 mice). Individual session data points/means (filled dots) are shown with population mean (squares), median (horizontal lines), quartiles (box) and extreme (vertical lines) values. **D**: Comparison of sample-wise spatial spread of pre- (black) and post-ablation (orange) normalized paths (thin traces: individual sessions, thick traces: means, 0: movement start, 1: movement end). Comparisons are between mean values binned in 5 consecutive path intervals. **E**: Left: Depiction of the shape-preserving alignment transform correcting for spatial dispersion for three example pre- and post-ablation trials. Right: 3D movement trajectories in C after spatial alignment. **F**: Spatial spread and stereotypy comparison of spatially aligned trajectories. n.s.: p>0.05, *: p<0.05, **: p<0.01 (linear mixed effects model fit).

We photoablated 14.7 ± 7 (mean ± s.d.) neurons across 6 mice and compared reach performance and execution before and after the manipulation. Mice continued to accurately grasp water droplets post-ablation, with no significant changes in the percentage of successful grasps or total movement path lengths (Fig. 2C). However, we noted a conspicuous modification of movement trajectories, which became markedly more stereotyped across trials. This increase in shape consistency is qualitatively apparent when comparing individual sessions (Video 2) and the one-dimensional position and velocity time series plots (Fig. S1A). To quantify this effect, we performed Principal Component Analysis (PCA) on the movement trajectories for each session and defined a stereotypy index as the cumulative variance explained by the first three principal components (see Methods). Following the ablation, we observed a significant increase in PCA-based stereotypy (Fig. 2C), indicating a reduction in the trial-to-trial variability of reaching paths.

Because this measure reflects both the geometric consistency and the spatial dispersion of reach trajectories, we next sought to isolate the individual contributions of these two factors. We first quantified the spatial dispersion of temporally normalized paths as the mean Euclidean distance to the centroid at each time sample. Although starting position variability remained the same, the paths became progressively more dispersed throughout the course of the reach after the ablation (Fig. 2D). Consequently, the observed post-ablation increase in PCA stereotypy occurred despite this heightened spatial dispersion, suggesting that it is primarily driven by more consistent trajectory shapes.

To directly compare variability in trajectory shapes, we used a shape-preserving alignment method that minimizes the confounding effects of spatial dispersion. Each trajectory was first re-parametrized by arc length, resampling the path at uniformly spaced points in distance traveled, so that they reflect equivalent fractional positions along the geometric path across trials. The resulting trajectories were then optimally superimposed on a template shape using generalized Procrustes analysis^39^. The two transforms thereby remove the main sources of non-shape variability (i.e. temporal non-uniformity, spatial translation and rotational shift) while leaving the geometric form of each trajectory intact (Fig. 2E, Fig. S1B). Following the correction, sample-wise spatial dispersion was strongly reduced and differences minimized while the PCA stereotypy remained significantly higher post-ablation (Fig. 2F). Together, these findings indicate that fS1 proprioceptive neurons are essential for constraining spatial dispersion while simultaneously preserving the natural shape variability of reaching trajectories.

Because ablated neurons were identified based on passive movements in the horizontal plane, they may represent a biased subpopulation preferentially controlling feedback of horizontal limb displacements. To test this, we analyzed trajectories separately in the horizontal and two vertical (coronal and sagittal) planes of motion. Significant changes were observed across all three dimensions (Fig. S2A–C), indicating that the ablated neurons contribute to feedback control of limb movements beyond the horizontal plane. We also show that the observed increase in stereotypy is robust to the choice of principle components used for analysis (Fig. S2D) and that the finding can be replicated with a different dimensionality reduction method (Supplementary Fig. 2F). Interestingly, the magnitude of the effect did not scale with the number of ablated neurons across individual mice (Fig. S2E). The lack of correlation suggests that the observed change in reach execution is not a simple linear function of cell loss but instead reflects a network-wide disruption.

Even a sparse ablation thus seems sufficient to disrupt the coordinated activity of a broader, functionally linked S1 population, consistent with previous findings^21^.

### Primary movement kinematics are largely unaffected by the fS1 cell ablation

To assess the impact of selective ablation on movement kinematics, we decomposed individual trajectories into reach, grasp, and retract components and analyzed the main sequence for each (Fig. 3A). We found that the main sequence slopes (MSS, linear relationship between amplitude and peak velocity) differed significantly across the three phases. Specifically, the MSS for grasping (11.1 ± 2.3, mean ± s.d.) was significantly greater than that of both reaching (2.0 ± 0.9; p<10^-16^) and retracting (2.3 ± 1.1; p<10^-18^). This indicates that peak velocity scales more steeply with movement amplitude during grasping than during the other phases. These kinematic distinctions suggest that the reach, grasp, and retract components may be governed by phase-specific neural controllers (e.g. with different forward gains), or by the optimization of distinct velocity costs at each stage of the movement.

**Figure 3.**
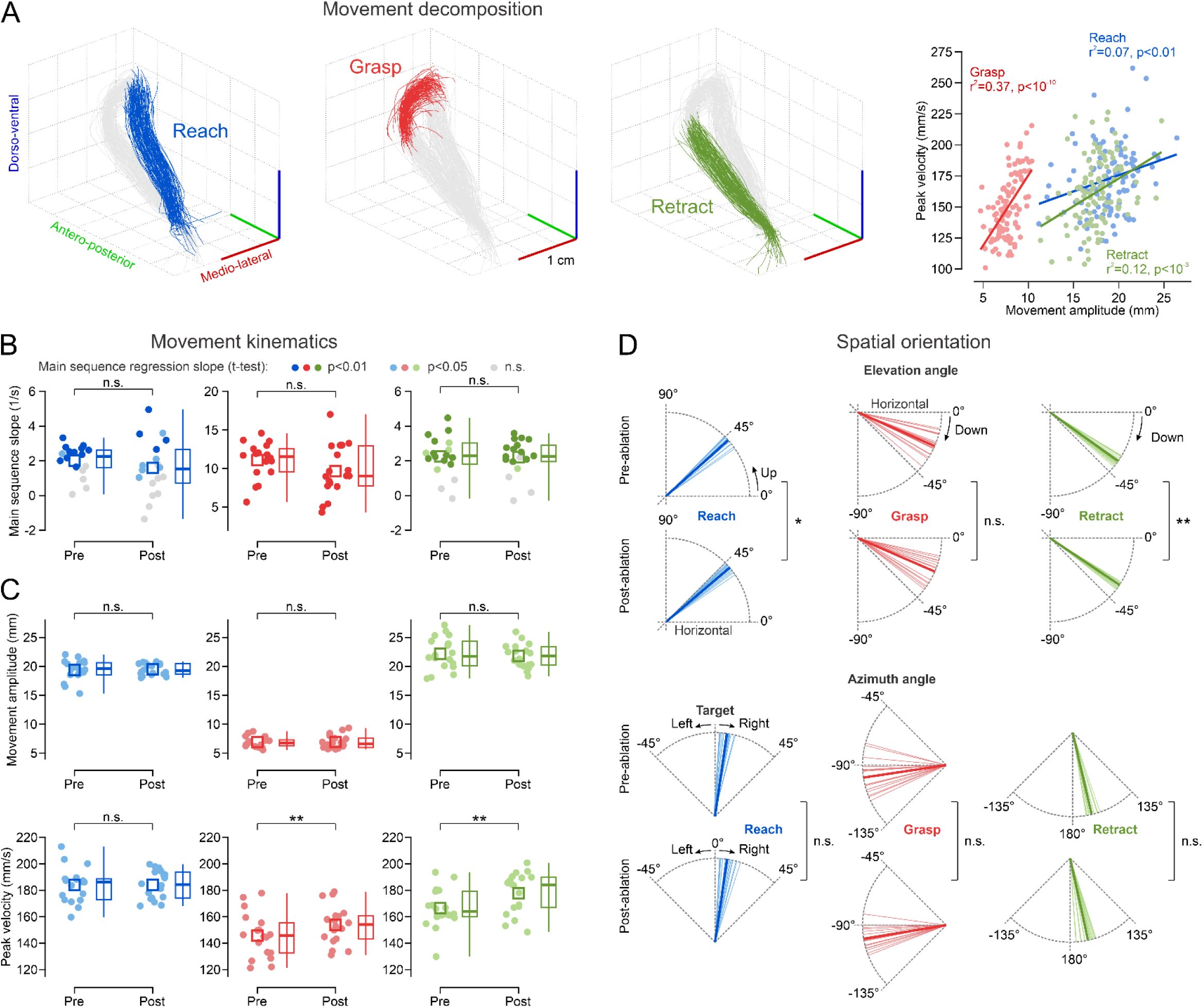
Changes in reaching kinematics after selective microablation of proprioceptive fS1 neurons. **A**: Trajectories of an example session decomposed into reach, grasp and retract components (all trials shown). Right: main sequences (with linear regression fits) depict how movement amplitude scales differently with peak velocity for the three components. **B, C**: Comparison of pre- and post-ablation main sequence slopes, amplitudes and peak velocities (symbols as in Fig. 2). Data point colors and p values in B correspond to the significance of the main sequence regression in individual sessions. **D**: Comparison of pre- and post-ablation azimuth and elevation angles (thin lines: session means, thick lines: population mean). n.s.: p>0.05, *: p<0.05, **: p<0.01 (linear mixed effects model fit).

Comparison of kinematics before and after ablation showed that the MSS remained the same across all movement components (Fig. 3B). No significant changes in movement amplitudes indicate an absence of dysmetria (Fig. 3C), though grasping and retraction were executed with higher peak velocities (Fig. 3C). To assess whether the ablation impaired spatial orientation or gravity compensation, we quantified changes in azimuth and elevation angles across movement components (Fig. 3D). Angle measurements were based on the dominant direction of motion calculated as the principal movement axis via PCA. We observed no significant directional shifts, except for decreased elevation angles during reach and retraction. The stability of these angular components indicates that the spatial representation of the workspace and the ability to calculate directional vectors remain robust.

Collectively, these results imply that the loss of fS1 proprioceptive neurons minimally disrupts motor commands governing movement kinematics or spatial control. Maintaining the natural variability of reaching paths therefore seems to be their primary function, which may rely on the optimal integration of noisy sensory feedback.

### OFC model simulations identify state estimation as the source of increased stereotypy and spatial dispersion

We investigated whether the optimal integration of sensory feedback could explain our findings by simulating the effects of cortical proprioceptive loss using an Optimal Feedback Control (OFC) model (Fig. 4A). OFC has served as a useful framework for motor control that captures many features of motor performance and neural processing^26,40–43^. Critically, because OFC optimizes control not for a desired trajectory but for task goals, it adjusts feedback gains only when variations in limb state interfere with those goals (i.e. the minimum intervention principle^26^). In the presence of sensory and system noise, such flexible adjustment of model parameters reproduces the trial-to-trial variability of movement paths observed in biological systems^26,43^.

**Figure 4.**
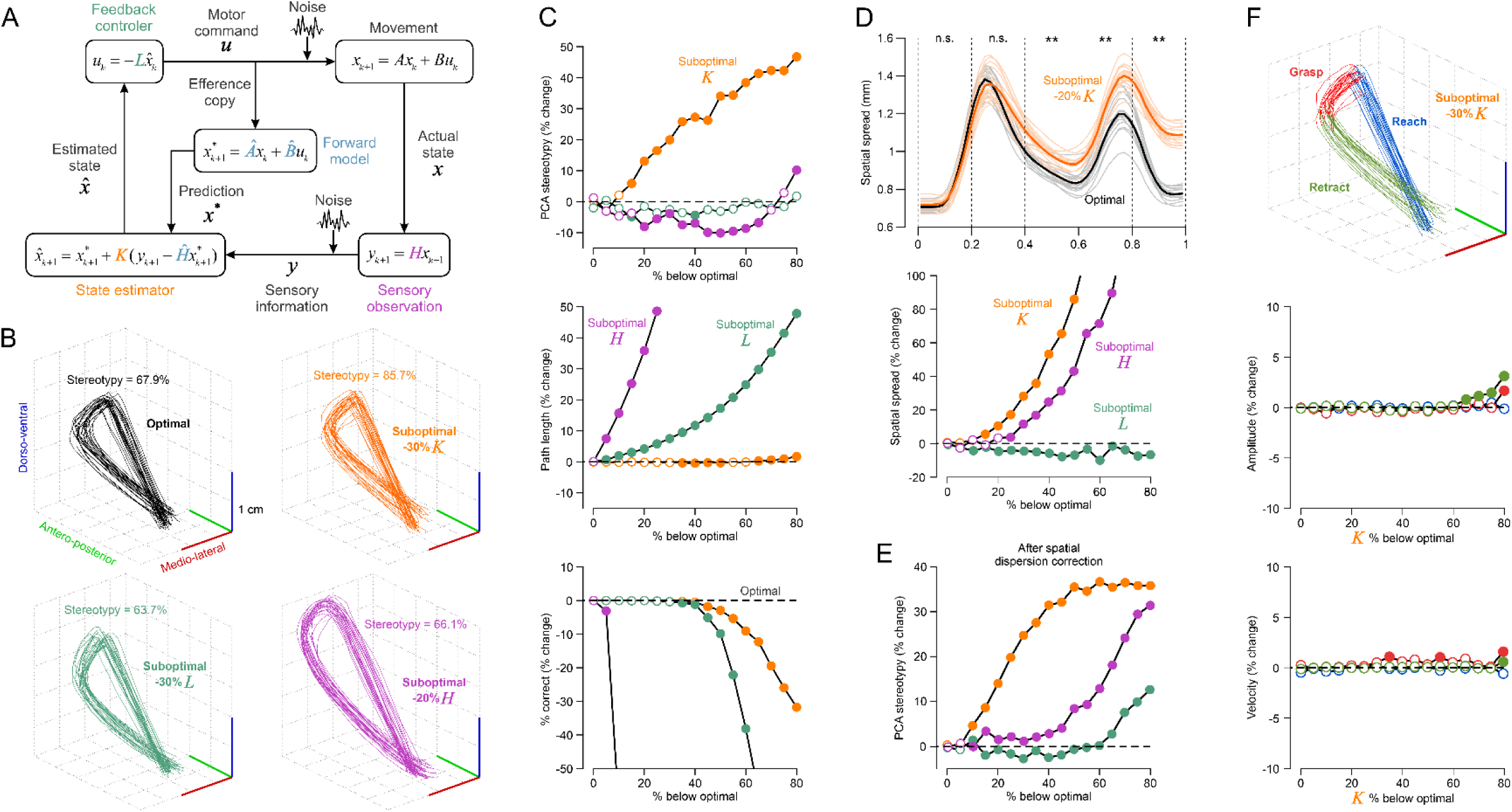
OFC model simulations. **A**: optimal feedback control model. **B**: Simulated movement trajectories (N=100 trials) with optimized parameters (black) and suboptimal (downscaled) K, L and H parameters (colors). The computed stereotypy index is provided for each simulation. **C**: Simulated changes in stereotypy, path length and correct performance with respect to optimized parameters (zero, black dotted line) for different levels of downscaled K, L and H parameters. Filled symbols denote significant differences from zero (p<0.01). **D**: sample wise spatial spread for simulated sessions (N=18) with optimal parameters and a 20% downscaled Kalman gain (data as in Fig. 2D). Bottom: Simulated changes in spatial spread (data as in C). **E**: Simulated changes in stereotypy after spatial trajectory alignment. **F**: Simulated trajectories decomposed into reach, grasp and retract components (top) and their changes in amplitude and peak velocity with respect to optimal parameters for different levels of K downscaling.

We extended the mathematical implementation of OFC^26,44^ to three dimensions to simulate trajectories of the reaching task. The simulation was based on minimizing a cost function that included an instantaneous position error cost during each of the three movement phases. Therefore, the goal of the model was to move the limb to a distinct spatial position during reach, grasp and retract periods in the presence of signal-dependent and additive sensory and system noise (see Methods for details). Using the computed optimal parameters, each simulated trajectory was unique because feedback corrections deviated the limb from a nominal path differently on each trial (Fig. 4B, black trace). This variability allowed us to compute a PCA-based stereotypy index identical to the one used for experimental data.

We simulated the effect of fS1 neuronal ablation by downscaling model parameters below their optimal values. We then quantified changes in the stereotypy index and task performance, including path length and the percentage of correct trials (Fig. 4B, C). Downscaling the gain of the feedback controller (*L* in Fig. 4A) or the sensory observation matrix (*H*) failed to increase stereotypy and instead produced hypermetric reaches that increased path lengths and errors. Modifying parameters of the internal forward model (*A*, *B* and *Ĥ* produced unstable trajectories (even for <20% suboptimal, data not shown) confirming the high reliance of OFC on an intact forward model^40^. In contrast, reducing the Kalman gain (*K*) in the state estimator reproduced the experimental effects of fS1 ablation: it increased path stereotypy while keeping path length and correct performance constant (until reducing the gain beyond ≈50%). This result remained robust across model initializations yielding different baseline stereotypy levels in the optimal condition (Fig. S3).

Comparing spatial variability of the simulated trajectories showed that a suboptimal K drove a progressive increase in sample-wise dispersion (Fig. 4D), replicating the patterns observed following selective proprioceptive fS1 neuron ablation (Fig. 2D). After correcting for spatial dispersion via resampling and alignment, increases in pure shape stereotypy also proved most sensitive to downscaling the K parameter (Fig. 4E). Furthermore, in line with experimental results, a suboptimal K left the amplitudes of decomposed reach, grasp and retract phases largely unaffected (Fig. 4F).

The reason that deactivating state estimation produces more stereotypical movements seems to lie in how the Kalman gain weighs information. The Kalman gain determines the relative contribution of the internal state prediction versus external sensory information to the feedback. Downscaled values signify a lower reliance on sensory signals^40^. It therefore seems that unique feedback corrections typically induced by noisy sensory signals are reduced compared to an optimized *K* parameter. A lowered Kalman gain, while increasing spatial dispersion due to suboptimal signal integration, prevents the movement path from deviating across multiple dimensions, causing the system to default more closely to a single nominal trajectory. The maintenance of normometric movement components (Fig. 3D) can be explained by the forward model internal feedback compensating for the decreased contribution of sensory information. This compensation is absent and results in dysmetria when simulating ablation of *H* and *L* parameters (Fig. 3B,C).

We conclude that the observed shift towards more spatially dispersed, normometric movements but with more stereotyped trajectory shapes (Fig. 2C, D) can be explained by the OFC model. Within this framework, fS1 proprioceptive neurons contribute primarily to optimizing state estimation (*K*) by integrating sensory feedback with internal predictions, rather than participating in direct feedback control (*L*) or sensory information coding (*H*).

### Temporal phasing of fS1 engagement in motor control

While these simulations highlight a specific role for the ablated neurons in state estimation, downscaling the Kalman gain failed to reproduce the increased peak velocities observed during the grasp and retract phases (Fig. 3C, 4F). This discrepancy could be explained by off-target effects of the ablation on the local network, such as the layer 5 corticospinal system or sensory to motor cortico-cortical circuit, suggesting that fS1 likely has functional roles broader than state estimation alone. We therefore moved from selective cell ablation to a complete lesion of fS1 to determine if the region contributes to additional aspects of motor execution.

We photo-ablated fS1 using targeted high-power infrared laser pulses^45^ delivered with a low-NA optical fiber through the cranial window (Fig. 5A, see Methods). We selectively ablated four sites covering the proprioceptive activation map in fS1^1^. Lesions were confirmed with pre- and post- two-photon imaging and histological verification (Fig. 5B,C).

**Figure 5.**
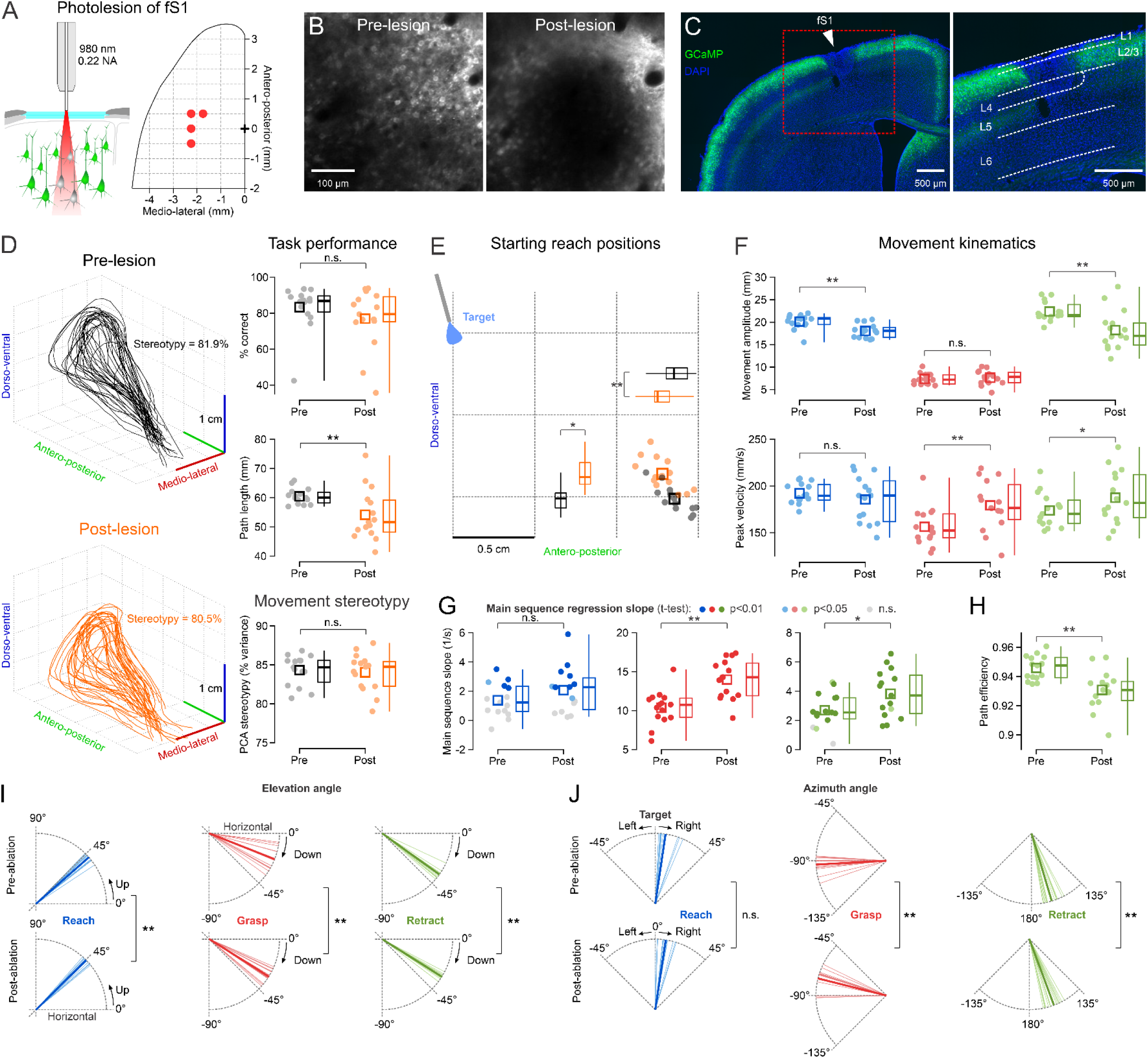
Changes in movement trajectories after complete fS1 lesion. **A**: Schematic of targeted cortical laser lesions (left) with coordinates of the four lesioned locations covering the fS1 proprioceptive map (right). **B**: Pre- and post-lesion two photon averaged images of a target location. **C**: Histological verification of the fS1 lesion. **D**: Pre- and post-lesion 3D movement trajectories of example sessions from the same mouse (left, only the first 25 trials are shown for clarity). Right: comparison of correct performance, trajectory path length and stereotypy pre- and post-lesion (N=15 sessions, 5 mice). Individual session data points/means (filled dots) are shown with population mean (squares), median (horizontal lines), quartiles (box) and extreme (vertical lines) values. **E**: Comparison of pre- and post-lesion starting reach positions in dorso-ventral and antero-posterior dimensions (all symbols are as in D). **F, G, H**: Comparison of pre- and post- lesion main sequence slopes, amplitudes, peak velocities and path efficiencies (symbols as in D). Data point colors and p values in G correspond to the significance of the main sequence regression in individual sessions. **I, J**: Comparison of pre- and post-lesion azimuth and elevation angles (thin lines: session means, thick lines: population mean). n.s.: p>0.05, *: p<0.05, **: p<0.01 (linear mixed effects model fit).

In contrast to the effects of selective neuronal loss, movement path lengths were significantly reduced, and the previously observed increase in PCA-based stereotypy was absent (Fig. 5D). This suggests a disruption of feedback processes that extends beyond simple state estimation (Fig. 4). Despite shorter trajectories, task proficiency (successful grasp %) remained stable (Fig. 5D); this was achieved through a compensatory strategy in which mice initiated reaches from starting positions significantly closer to the target in both dorso-ventral and antero-posterior axes (Fig. 5E).

Analysis of individual movement components confirmed the emergence of hypometric movements during the reaching and retraction phases, while grasping amplitude remained stable (Fig. 5F). Despite the reduction in reach distance, we again observed significantly higher peak velocities during grasping and retraction only (Fig. 5F). Notably, this was accompanied by an increase in their main sequence slopes (Fig. 5G).

Beyond kinematic changes, we examined whether fS1 lesion modifies the spatial topology of the movement by quantifying changes in azimuth and elevation angles, as well as path efficiency (the ratio of displacement amplitude to path length). While an efficiency value of 1 represents a perfectly straight trajectory, lower values indicate increasingly curved or inefficient movements. We observed a significant reduction in path efficiency during the retraction phase only (Fig. 5H), whereas reach and grasp components remained at pre-lesion efficiency levels (data not shown). Correspondingly, significant shifts in both elevation and azimuth angles emerged mainly during the grasp and retract phases, while the reaching movement remained more spatially stable (Fig. 5I,J).

Considered alongside the component-specific changes in peak velocity and main sequence (Fig. 5F,G), these results indicate that while the initial hypometric reach remains spatially targeted with consistent kinematic scaling, the subsequent grasping and retraction phases exhibit significant spatial and kinematic instability. This implies that fS1-mediated feedback control is temporally phased. It is engaged late in a movement sequence as control transitions from an initial, preprogrammed, subcortically driven^46–48^ and possibly open loop^49,50^ motor launch to cortical refinement of grasp and return phases. Such a framework aligns with the late-phase online adjustment of reach kinematics^51^, supporting a model where cortical feedback starts rectifying the trajectory as the limb approaches the target.

### Phase-specific classification of motor deficits

Although these discrete kinematic changes support a temporally phased role for fS1, they do not capture the full, potentially nonlinear impact of the lesion on movement dynamics. More subtle disruptions in temporal dependencies might already exist from the onset of reaching, allowing for a level of discriminability comparable to that in later grasp and retract phases. To test this, we employed a Long Short-Term Memory (LSTM) neural network, a model specifically designed to extract features from sequential data (Fig. 6A). By training the model to classify trajectories as pre- or post-lesion, we could assess whether motor disruptions are equally identifiable during phases where traditional kinematic measures show less significant change.

**Figure 6.**
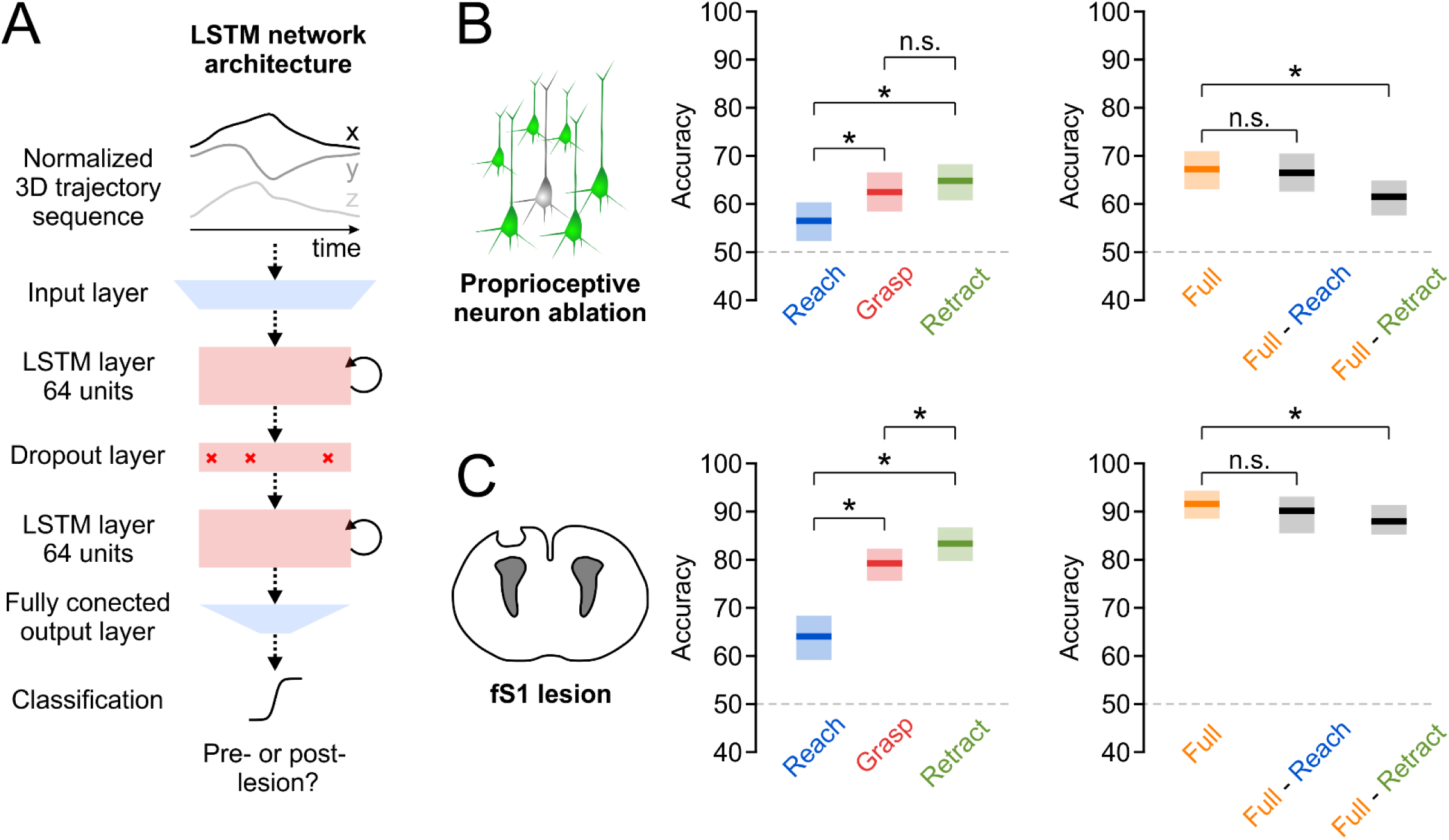
Machine learning classification of movement phases. **A**: Architecture of the neural network classifier. **B**: Comparison of pre-vs. post-ablation classification accuracy for the proprioceptive neuron microablation experiment. Mean (± 95% confidence intervals) classification accuracy is shown for the three movement phases as well as for the full and truncated trajectories. **C**: Analogous data for the broad fS1 lesion experiment.

To focus the network on movement dynamics, all trajectories were normalized to a range of −1 to 1 prior to training. This step ensures that classification is driven by disturbances in spatiotemporal patterns and shape rather than spatial location and scale. Furthermore, each movement phase (reach, grasp, retract) was rescaled independently. This mathematical decoupling prevents spatial offsets at the end of one phase from propagating into the next, forcing the network to classify trials based on phase-specific movement properties rather than cumulative spatial biases.

We trained and evaluated separate LSTM models on the three trajectory components, repeating the procedure 199 times with independent random dataset splits to generate a distribution of classification accuracy. Classification was considered above chance if the 2.5th percentile of the accuracy distribution exceeded 50%. Accuracy was deemed significantly different between conditions if the mean of one distribution fell outside the 95% confidence interval of the other.

Above chance classification was achieved in all three phases, successfully distinguishing pre- and post- trajectories for both selective proprioceptive ablations (Fig. 6B, left panel) and full fS1 lesions (Fig. 6C, left panel). However, consistent with our discrete kinematic findings, the LSTM classifier achieved higher accuracy during the grasp and retract phases compared to the initial reaching phase across both experimental groups. To quantify the relative contribution of early vs. late components to the classification of the entire movement, we compared model performance on full trajectories against truncated sequences where either the initial reach or the final retraction component was removed (Fig. 6B,C, right panels). In this comparison, we found that the reach contributed negligibly as its removal did not lower classification accuracy. In contrast, the exclusion of the retraction significantly impaired performance, highlighting its critical contribution to the classifier.

We conclude that while indicators of cortical proprioceptive loss are detectable during the initial reach, this early phase is less essential for capturing the resulting motor impairments. fS1-mediated feedback control seems to be primarily engaged during the subsequent grasping and retraction aspects of movement, which provide the most discriminative features for identifying movement deficits.

## Discussion

### The advantage of selective ablation over neural encoding analysis

We examined the function of fS1 proprioceptive neurons, identified by their responses to passive limb displacements, within the feedback control of reaching movements. We moved beyond standard encoding analyses that correlate neural activity with motor variables. Although such analyses can distinguish between signals representing low-level joint kinematics versus high-level task parameters^52^, they offer limited insight into the actual function of these neurons. Indeed, encoding studies often give the broad conclusion that sensorimotor cortical activity covaries with a vast array of kinematic signals alongside other complex behavioral features^52–56^, highlighting the difficulty in assigning precise causal roles. This overlap arises because functional ensembles, whether specifying motor commands, controlling feedback, or integrating predictions, all contribute to the same task goals and interact with the same peripheral plant. As their activity necessarily reflects these identical physical coordinates, it is expected that similar heterogenous neural codes would appear throughout the sensorimotor system. Critically, none of these signals need to be explicitly specified by the neuronal ensembles; they might emerge as a natural consequence of being embedded within a feedback controller that optimizes motor commands based on the state of the effector and task goals^43,57^. They arise because the system is reacting to the physical reality of the limb and its environment when working towards the goal. Relying on such correlations can thus lead to erroneous conclusions regarding the actual causal role of the underlying neurons. To bypass these ambiguities and isolate the function of the proprioceptive fS1 ensemble, we employed targeted microablations to directly observe the impact of their loss on reaching behavior.

### L2/3 neurons as optimal state estimators

Targeted manipulation of a small number of functionally labeled neurons in sensory cortices has been shown to bias perception^19,20,22–24,58^. Here, we demonstrate that selective sensory neuron removal also modifies motor control. Our unexpected finding was that movement trajectories became more dispersed but also more stereotyped (Figure 2). This modification suggests a seemingly paradoxical dual role for fS1 proprioceptive L2/3 neurons: constrain spatial dispersion while maintaining shape variability of movement paths. OFC model simulations explain away this paradox by showing that both effects can emerge from a suboptimal weighting between sensory feedback and internal predictions (Figure 4), implying that the underlying neural network participates in state estimation rather than merely encoding peripheral signals. This integrative role reinforces the view that S1 is not a passive sensory map, but an active component of an internal body (or limb) model^59,60^. At the physiological level, L2/3 cortical neurons likely integrate bottom-up thalamic inputs with top-down predictions from deeper layers^61–63^, functioning as a general predictive processing node in sensory cortices, that supports both perceptual mismatch signaling^60,64^ and, as shown here, motor command optimization.

It should be noted that a reduced Kalman gain can also be produced by increasing sensory noise. Instead of manually lowering the gain, one could recompute an optimal *K* based on a noisier sensory signal. The resulting lower gain would naturally increase the weighting of internal predictions, effectively replicating the effects of the ablation. This alternative implies that the ablated neurons act to reduce noise in the feedback signal upstream of the state estimator, rather than being part of the estimator itself. However, in this scenario the controller parameters must be re-estimated prior to simulating the new trajectories, which assumes that rapid plasticity occurs immediately after the ablation. In contrast, simply downscaling a model parameter requires no such reorganization of the underling neural circuits. It is also unclear how cortical neurons could actively reduce noise from sensory afferents without integrating an additional source of information. Therefore, a direct reduction of a previously optimized Kalman gain provides a more parsimonious account of our results.

Consistent with our findings, the effects of inactivating the parietal somatosensory area 5 in the primate cortex was most consistent with downscaling the *K* parameter in OFC model simulations of corrective limb movements^40^. Although the authors of that study did not analyze stereotypy, their results also show that both manipulations primarily increase spatial dispersion (i.e. endpoint variability) while minimally affecting movement kinematics. These parallels suggest that state estimation is distributed across both primary and higher order somatosensory cortical areas. Alternatively, these areas may be less anatomically segregated in the mouse brain. The mouse S1 may thus serve as a more diverse integrative hub, functionally resembling Brodmann’s areas 2 and 5 more closely than the primary sensory area 3.

### Loss of specificity with broad cortical lesions

Broad lesion of the proprioceptive map^1^ in fS1 masked the effect of the selective proprioceptive ensemble removal. Instead, it produced modifications in movement kinematics that cannot be directly related to the proposed role of fS1 neurons in optimal state estimation. This highlights the fundamental principle that deciphering functional roles of neuronal ensembles within a local cortical network requires fine-grained, cellular-level causal perturbations.

Importantly, the broad lesions did not cause a complete breakdown of motor control, but rather an impaired movement scaling and a re-adjustment of the main sequence relationship (Figure 4E-G). The fS1 feedback controller is therefore most likely nested within recurrent sensorimotor loops, occupying a hierarchical position above subcortical control pathways. In this role, it acts to extend movement trajectories and downregulate the speed to amplitude sensitivity, effectively refining a high-gain, unscaled subcortical drive.

Additionally, the lesion modified the spatial topology and trajectory efficiency of the movements (Figure 4H-I). We abstained from attempting to simulate such widespread effects with the OFC model because the extensive damage does not affect a specific computational unit of the cortical circuit. In addition to the complete removal of local circuitry it inevitably disrupts distant networks targeted by S1 projections in other cortical and subcortical structures. While the OFC model can effectively simulate isolated changes of a single feedback loop, it is less suited for characterizing lesions that affect multiple aspects of a hierarchical control system. More physiologically relevant hierarchical feedback controllers^65^ can better interpret how disruption of the entire S1 affects movements. Furthermore, the high number of altered variables creates multiple modeling solutions and tends to make the simulation independent of the optimal regime the OFC model is intended to describe.

### Hierarchical control of movement phases

Comparing the effects of both selective and broad lesions across sequential movement components revealed that they disrupt late-stage grasping and retraction significantly more than initial reaching (Figures 5 and 6). This implies that fS1-mediated feedback control is temporally phased during reach-to-grasp behavior. Our results align with the late-phase online adjustment of reach kinematics^51^, supporting a model where cortical feedback starts rectifying the trajectory as the limb approaches the target.

The late fS1 engagement suggests that control transitions, throughout the movement sequence, from an initial, preprogrammed, subcortically driven^46–48^ and possibly open loop^49,50^ motor launch to cortical refinement of grasp and return phases. The transcortical loop is thus nested within a hierarchical motor network including multiple recurrent controllers^65^. Under this framework, subcortical circuits might orchestrate reaching independently of, and prior to, fS1 involvement, allowing the cortex to remain unencumbered by low-level control details. Furthermore, grasping and retraction involve prehension and retrieval of the water droplet, requiring finer motor precision than simple arm extension. A subcortical network responding to an initial cortical pulse may lack the computational capacity to execute movement phases containing these fine components without the integration of transcortical feedback. This provides a mechanistic explanation for why optogenetic silencing of fS1 fails to impair motor performance when restricted to the reach phase^34^ or applied during forelimb movements lacking high precision requirements^13^.

An alternative possibility is that fS1 supports a modular control network, where sequential phases of complex movement are governed by distinct groups of cortico-spinal neurons (CSNs) in caudal and rostral forelimb motor areas^66^. In this scenario, CSN subsets mediating grasping and retraction may receive stronger anatomical or synaptic fS1 drive compared to those mediating reaching. The differential drive may also result in different noise statistics across the three components.

Finally, our previous work demonstrated that body-petal directions are overrepresented in the preferred tuning of proprioceptive fS1 neurons during passive limb displacements^1^. This anisotropy may reflect a functional, experience-dependent specialization of fS1, optimized for feedback control of movements involving the retrieval of a grasped object toward the body.

### Limitations

Our results are based exclusively on paw trajectories. Tracking the full pose of the mouse forelimb would provide a more complete picture by allowing us to compare the effect of ablations in both endpoint and joint angle spaces. Optimal control predicts that variability is lower in the task-relevant endpoint position than in the task-irrelevant joint angles^67,68^. While we might therefore expect changes in movement stereotypy to manifest differently across these two spaces, testing this hypothesis remains challenging. Specifically, video-based tracking of mouse forelimb joints is highly imprecise because proximal joints lack clear visual features and do not move in conjunction with the skin^1^. Moreover, the OFC simulations are based on moving a point mass which does not represent the limb’s mechanical properties. The model therefore does not simulate the true biomechanical complexity of the movement. It simply tests whether the nervous system employs a solution consistent with the principles of optimal feedback control. Taken together, practical solutions for joint tracking and accounting for the physics of limb movement, combined with selective neuronal ablation experiments, can further extend the findings of the current study, thereby revealing how unique computational aspects of motor control are implemented in the mouse brain.

## Acknowledgements

This study was supported by the Swiss National Science Foundation (Grant TMCG-3_213672).

## Methods

### Animals

All experiments were performed with adult double transgenic Rasgrf2-Cre x Ai148 mice (Jackson laboratory, #022864, Jackson laboratory #030328; 17 males, 8 to 12 weeks at the start of the experiments) expressing Tet-controllable GCaMP6f in layer 2/3 cortical neurons. Mice were housed in groups of five adults maximum per cage and kept in an animal facility under standard housing conditions (12h light/dark cycle) outside experimental hours. All procedures were approved by the Fribourg Cantonal Commission for Animal Experimentation and in accordance with the veterinary guidelines.

### Surgical procedures

Surgeries for chronic calcium imaging experiments were performed as previously described^1^. Briefly, mice were anesthetized, the scalp was exposed, and a titanium head frame was fixed to the skull. A craniotomy was centered over the left forelimb somatosensory cortex (fS1) and covered with a hand-cut cranial window, allowing chronic optical access for 2-photon imaging. Following recovery, mice were injected intraperitoneally with trimethoprim (Sigma-Aldrich T7883; at 0.25mg/g of body weight per day over 3 consecutive days) to induce Cre recombinase dependent expression of GCaMP6f. Trimethoprim was dissolved in DMSO (Sigma-Aldrich #34869; stock solution, 100 mg/ml) and further diluted in saline prior to injection (62.6mg/ml final concentration). Behavioral experiments began 2 weeks after surgery.

### Single cell microablation

Proprioceptive neurons were first identified based on their significant responses to passive limb displacement. Targeted photoablation was then performed by scanning the somata with 920 nm femtosecond laser pulses (<100 fs) at 500 mW for a maximum of 10 s in awake mice. Scanning was immediately terminated upon the detection of a somatic calcium fluorescence increase. Successful ablation was verified 24 hours post-procedure, confirmed by either the total disappearance of the targeted cell or the absence of both spontaneous and stimulus-evoked calcium transients.

### Cortical lesion

Focal cortical lesions were induced in anesthetized mice using a 200 μm (0.22 NA) optical fiber positioned via a stereotaxic micromanipulator over the cranial window. We targeted four locations covering the proprioceptive activation map in fS1 (ML, AP in μm: −2250, –500; –2250, 0; –2250, +500; and –1750, +500). At each site, five 1 s pulses of continuous wave 980 nm laser light (power >1 W) were delivered with a 10 s inter-pulse interval.

### Histology

Mice were transcardially perfused with 4% paraformaldehyde (PFA; in 0.1M sodium phosphate buffer, pH 7.4). The brains were dissected and post-fixed overnight in the same solution, then rinsed (3×10 min) with 0.1M phosphate buffered saline (PBS). The tissue was cryoprotected at 4°C in 15% and 30% sucrose (in PBS) for 24 h each. Brains were cut in coronal sections (40 µm thick) using a cryostat (HM535 NX, Thermo Scientific) and mounted on Superfrost Plus Adhesion slides (Fisher Scientific AG, Reinach, Switzerland). Sections were then stained with DAPI (1:1000 in PBS; Sigma-Aldrich, #32670-F) for 5 min, followed by three PBS washes (5 min each), and mounted on coverslips with fluorescent medium (Dako, Agilent Technologies, #S3023). Images were acquired with a wide-field microscope (Leica DM6B Navigator, Leica, Germany) and processed with LAS X software. Post-acquisition image processing was performed using ImageJ/Fiji (NIH; RRID: SCR_002285).

### Behavioral procedures

All behavior was controlled and measured with real-time protocols running on the Bpod State Machine system (Sanworks).

#### Passive forelimb movement task

Passive limb displacement was performed in head-fixed, awake mice using a custom robotic manipulandum, following a previously described protocol^1^. Briefly, mice were trained to grasp and maintain contact with the manipulandum endpoint while their right limb was displaced from a central ’home’ position to one of eight co-planar peripheral targets. Each centrifugal movement was followed by a centripetal return to the home position. Movements spanned 5 mm and followed a trapezoidal velocity profile (3 cm/s). To ensure continuous contact, a capacitive sensor (MPR121, Adafruit) was used to detect limb releases. Any such release resulted in an immediately aborted trial. Mice were rewarded with a water droplet only upon completion of a full trial (both centrifugal and centripetal phases) without release. Only neuronal responses from these successful trials were included in the subsequent analysis.

#### Reach-to-grasp task

Unrestrained mice were placed in a transparent acrylic chamber (100 by 100 by 120 mm). A vertical slit (20 by 10 mm) centered on one of the vertical walls at a height of 60 mm allowed for arm extension outside the chamber. Trials were initiated with a 300 ms auditory cue and the simultaneous delivery of a water droplet via a 1.3 mm diameter tube, controlled by a miniature solenoid valve (The Lee Company, LHDA1231215H). The reward was positioned 3 mm above, 15 mm anterior, and −3 mm lateral to the slit center to specifically elicit right arm reaching. Arm extensions outside the chamber were detected using an infrared fiber-optic sensor (Panasonic FX-301, FT-31 fiber), which provided the timestamps for video paw tracking alignment and analysis. Reward contact was registered using a capacitive sensor (Adafruit, MPR121). To ensure data independence and isolate discrete reaching movements, analysis was restricted to the initial reach of each trial, provided no subsequent extensions occurred within a 350 ms window. New trials were initiated only after an idle period of 2 s without further arm extensions. Data acquisition began once mice achieved baseline performance, characterized by consistent reaching and inter-trial intervals under 10 s. Recordings consisted of three pre-lesion and three post-lesion sessions conducted over consecutive days for each mouse.

Data acquisition, processing and analysis

#### Two-photon data acquisition and processing

Ca^2+^ imaging in fS1 was performed with a custom built two-photon microscope based on an open-source design (MIMMS 2.0, janelia.org/open-science) and controlled with Scanimage 5.7_R1 software (Vidrio Technologies) and National Instrument acquisition hardware. The ALCOR 920-2 fiber laser (Spark Lasers) at 920 nm was used for GCaMP two-photon excitation. The laser was focused with a 16x, 0.8 NA objective (Nikon), modulated with Pockels cell (350-80-LA-02, Conoptics) and calibrated with a Ge photodetector (DET50B2, Thorlabs). The mouse cortex was scanned with a frame rate of approximately 30 frames/s using a resonant-galvo scanning system (CRS8K/6215H, CRS/671-HP Cambridge Technologies). Emitted fluorescence was detected with GaAsP photomultiplier tubes (PMT2101, Thorlabs) and the acquired 512 x 512 pixel images were written to disk in 16 bit format.

Motion artifacts were corrected using a custom MATLAB script based on a rigid registration algorithm. Each acquired frame was aligned to an average baseline template. To determine the spatial offset, we computed the cross-correlation between each frame and the template in the frequency domain. Specifically, the two-dimensional discrete Fourier transform (DFT) of a frame was multiplied by the complex conjugate of the template’s DFT. The inverse Fourier transform of this product yielded the cross-correlation map. The coordinates of the peak value provided the vertical and horizontal pixel shifts required for alignment. To minimize edge artifacts, 10% of each image was cropped from the boundaries prior to computation.

Using the session mean and variance images, centers of active neurons with clearly identifiable morphologies were manually initialized. Regions of interest of individual neurons and background were then identified as spatial footprints using the constrained nonnegative matrix factorization method^69^ from the CaImAn MATLAB toolbox (github.com/flatironinstitute/CaImAn-MATLAB). The time-varying fluorescence activity of each neuron and its time-varying baseline fluorescence was extracted from the acquired images and used to compute Δf/f_0_ traces for neural activity analysis.

Proprioceptive neurons were identified based on their responsiveness to passive limb displacement stimuli. The stimulus evoked Δf/f_0_ fluorescence response was calculated as the difference between the peak post-stimulus value (0 to 1.5 s) and the mean pre-stimulus baseline (−0.75 to 0 s). To determine significance, we employed a randomization test (p<0.01) for each neuron. A null distribution was generated by re-calculating the response 1999 times using randomly shifted stimulus onset times across the session’s activity trace. An observed response was deemed significant if it exceeded the 99th percentile of this bootstrapped null distribution.

Directional selectivity was tested by fitting a Von Mises distribution to the neuron’s stimulus-evoked responses in the eight tested directions (*Circular Statistics Toolbox*^70^, MATLAB). Neurons with significant fits (p<0.05) were deemed directionally selective. The neuron’s preferred direction and directional selectivity were defined by the fitted μ and κ parameters, respectively.

#### 3D reach tracking

To track 3D reach trajectories, we utilized a stereo camera system consisting of two Basler dart daA720-520um cameras (640×480 pixels, 100 fps, external triggering). For each trial, we analyzed video segments spanning ±350 ms relative to the online reach detection timestamp.

Paw tracking was performed using a pretrained HRNet deep learning network (Matlab Deep Learning Toolbox, R2024b), which was adapted via transfer learning. We initially trained the network on a set of 2 300 manually annotated keypoints centered on the mouse right paw. This preliminary model was then used to predict paw locations across a broader dataset of 162 000 frames. After manual verification and correction of detection errors, this dataset was used to train the final model. This optimized HRNet detector was then deployed for automated paw tracking in all subsequent experiments.

Stereo camera calibration parameters were determined using the Matlab *Stereo Camera Calibrator App*. We then reconstructed the 3D coordinates of the tracked paw keypoints from the 2D stereo pairs using the *triangulate* function. To map these data from the camera-centric frame to world coordinates, we calculated a 3D affine geometric transformation. This was achieved by estimating the relationship between 15 corresponding pairs of calibration points across both coordinate frames via the *estimateGeometricTransform3D* function. Finally, we applied the resulting transformation to the tracked 3D paw positions in each frame using *transformPointsForward*, aligning all trajectories to a world coordinate system defined by the mouse’s antero-posterior and medio-lateral axes relative to the reach target.

#### Reach data analysis

Mice performed 95.7 ± 30.5 (mean ± s.d.) reach-to-grasp movements per session. We applied a second order, 9 frame long Savitzky-Golay filter to the raw 3D paw positions. To determine reach onset, we first identified the peak extension velocity along the anteroposterior axis (x) and moved backward in time until the velocity dropped below 20 mm/s. Similarly, the end of retraction was defined by moving forward from the point of minimum x-velocity until it exceeded −20 mm/s. Grasp onset and offset were determined using the peak velocity along the mediolateral (y) axis, moving backward or forward in time, respectively, until the velocity crossed a threshold of ±20 mm/s. The grasp onset/offset timings were used for reach end and retraction start, respectively. Correct trials were identified based on reward contact detection.

To quantify PCA-based stereotypy, we first standardized the length of all session trajectories through interpolation (normalized paths) and concatenated the three spatial dimensions. Principal component analysis (Matlab’s *pca.m* function) was then performed on the concatenated data. Stereotypy was defined as the cumulative percentage of total variance explained by the first three principal components. Sample wise spatial dispersion was calculated as the Euclidean distance to the centroid at each time sample of normalized paths. Statistical comparisons were made between mean values of binned data by dividing the paths into five intervals.

To isolate variability in trajectory shape from variability arising from spatial dispersion, the normalized paths were first re-parametrized by arc length, resampling the path at points uniformly spaced in distance travelled. The trajectories were then processed with Generalized Procrustes Analysis^39^ (GPA, MATLAB’s *procrustes* function), which iteratively estimates a mean template shape and computes, for each trajectory, the optimal rigid-body transformation (comprising of translation and rotation, but excluding scaling and reflection) that minimizes the residual sum of squares between each trajectory and the template. GPA was initialized with the medoid trajectory shape (the trajectory of the trial with smallest Procrustes distance to trajectories of all other trials).

We applied factor analysis (FA) as a complementary measure of stereotypy to PCA. FA partitions variance into a shared structured component (variance captured by common factors, termed FA communality) and a unique noise component (variance idiosyncratic to individual features, termed FA uniqueness). We used five factors for the analysis and only FA communality was counted as stereotypy. FA therefore provides a more conservative measure of stereotypy compared to PCA.

Movement amplitude of each component was defined as the 3D Eucledian distance between the start and end points. Peak velocity was similarly calculated as the maximum magnitude of the 3D velocity vector.

To determine movement orientation, we calculated the azimuth and elevation angles for each trajectory component. First, we identified the principal movement axis by performing PCA on the 3D spatial coordinates (treating dimensions as variables and time points as observations). This axis was defined by the first principal component, the eigenvector capturing the direction of maximum variance. We then converted this principal axis vector into spherical coordinates using MATLAB’s *vectorToSpherical* function to extract the azimuth and elevation. Session-wide angular means were computed using circular statistics (*Circular Statistics Toolbox*^70^, MATLAB).

Path efficiency was calculated for each trial as the ratio of displacement amplitude (Euclidean distance between 3D start and endpoint coordinates) to path length (sum of distances between all successive sample points of the trajectory).

Three sessions per condition were included for each mouse and when comparing all pre-and post-lesion behavioral metrics.

#### OFC model implementation

We modeled the translation of a point of mass *m* in three dimensions, driven by first-order actuator dynamics, following previous implementations of OFC models^26,44^. The discrete time dynamics of the control system were described as:

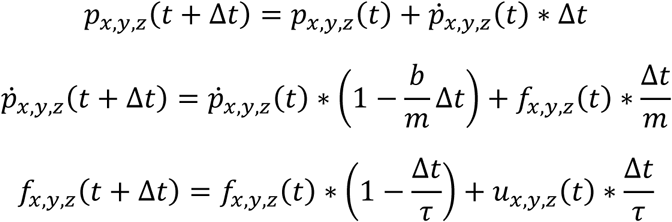

Where 𝑝(𝑡) and *ṗ*(𝑡) are the 3D position and velocity of the point mass as a function of time *t*, Δ𝑡 is the discrete time step, *b* is the damping constant, 𝑓(𝑡) are the forces produced along the 3 axes, 𝑢(𝑡) is the motor command and 𝜏𝜏 the time constant of the first order system.

The 3D point mass was initialized at position [𝑥_0_, 𝑦_0_, 𝑧_0_] and the control action was prescribed to transfer it to the first target [𝑥_𝑡1_, 𝑦_𝑡1_, 𝑧_𝑡1_] at time *t_1_*, then to the second target [𝑥_𝑡2_, 𝑦_𝑡2_, 𝑧_𝑡2_] at time *t_2_* and finally to target [𝑥_𝑇_, 𝑦_𝑇_, 𝑧_𝑇_] at the final time *T*. The target coordinates and sample times were chosen to simulate typical trajectories of grasp, reach and retract components observed in experimental data.

The discrete system state at sample *k* was thus described by the following column vector:

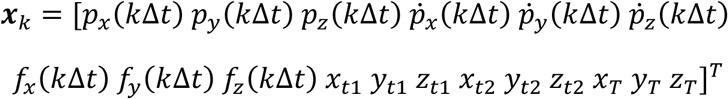

The initial state 𝒙𝒙_1_ was perturbed with the addition of Gaussian noise terms 𝒩𝒩(0, 𝜎𝜎_𝑖𝑖𝑖𝑖𝑖𝑖𝑡𝑡_), the estimated state was initialized as *x̂*_1_ = 𝒙𝒙_1_ and the command signal updated according to:

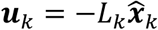

where 𝐿_𝑘_ is the optimal feedback gain at sample *k*.

The state vector dynamics and motor noise disturbances were described by:

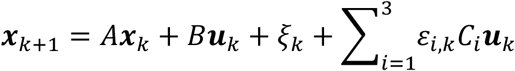

where *A* and *B* are matrices that describe the control system’s dynamics. The additive motor noise terms 𝜉_𝑘_are independent multivariate Gaussian variables with zero mean and covariance matrix 𝛺_𝜉_∈ ℝ^18×18^. 𝐶_𝑖_ are scaling factors that introduce multiplicative motor noise with 𝜀_𝑖,𝑘_ being Gaussian variables with zero mean and unity variance. For the additive noise to affect the motor commands only, we set the covariance matrix to

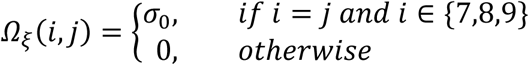

We set the scaling factors 𝐶_𝑖_ to:

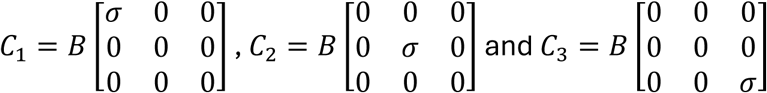

The noisy sensory feedback carries position, velocity and force information, and is updated at each sample according to:

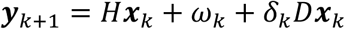

where H is the observation matrix:

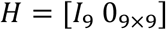

The additive sensory noise terms 𝜔_𝑘_ are independent zero mean Gaussians with standard deviations 𝜎_𝑠0_. Multiplicative sensory noise is introduced by scaling zero mean Gaussian variables 𝛿_𝑘_ with the factor 𝐷𝐷. We set the scaling factor 𝐷 = 𝜎_𝑠_[0.01𝐼𝐼_3_ 0.1𝐼𝐼_3_ 0_12×12_] for the multiplicative noise to affect position and velocity feedback only. A sensory delay *θ* was added to the feedback according to the system augmentation technique described in previous implementations^26,44^.

The internal prediction of the next state is computed as:

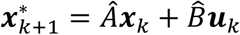

where 𝐴^ and 𝐵𝐵 represent the internal forward model of the system dynamics. The state estimate is then updated according to the Kalman filter:

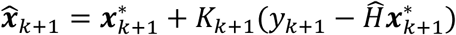

where the matrix 𝐾_𝑘+1_ is the optimal Kalman gain at sample *k+1* and *Ĥ* is in the internal model of the observation matrix. These internal processes are modeled as noiseless as adding prediction noise, which can be reasonably assumed to be orders of magnitude smaller than motor or sensory noise, had no significant impact on the results.

The optimal feedback controller and state estimator gains *L* and *K* were computed based on a modified Linear-Quadratic-Gaussian system solution^44,71^ and its implementation in Matlab (github.com/TomohikoTakei/kalman_lqg_pertpost1dof) that we adapted for three-dimensional trajectories. The optimization problem was to minimize the total cost function defined as:

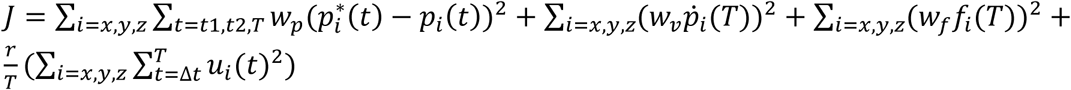

where the first term corresponds to reaching the three targets 𝑝𝑝^∗^ at times *t_1_*, *t_2_* and *T*, the second and third terms correspond to ending the movement with zero velocity and force at time *T* and the last term is the cost of control or effort penalty. The scaling weights 𝑒_𝑝𝑝_, 𝑒𝑒_𝑣𝑣_, 𝑒𝑒_𝑓𝑓_ and *r* define the relative Importance of the cost terms.

All optimizations and trajectory simulations were performed with the following parameter set: Δ𝑡 = 0.01, 𝑚 = 1, 𝑏 = 0.1, 𝜏 = 0.03, [𝑥_0_, 𝑦_0_, 𝑧_0_] = [0,0,0], [𝑥_𝑡1_, 𝑦_𝑡1_, 𝑧_𝑡1_] = [1.6,0.25,1.3], [𝑥_𝑡2_, 𝑦_𝑡2_, 𝑧_𝑡2_] = [1.6, −0.5,1.1], [𝑥_𝑇_, 𝑦_𝑇_, 𝑧_𝑇_] = [0,0,0.2], 𝑡_1_ = 0.2 𝑒, 𝑡_2_ = 𝑡_1_ + 0.08 𝑒, 𝑇 = 𝑡_1_ + 𝑡_2_ + 0.25 𝑒, 𝜎^2^ = 0.002, 𝜎_0_ = 0.01, 𝜎 = 0.1, 𝜎_𝑠0_ = 0.5, 𝜎_𝑠_ = 0.3, 𝜃 = 0.04 𝑒, 𝑒_𝑝_ = 1, 𝑒_𝑣_ = 0.1, 𝑒_𝑓_ = 0.01, 𝑒 = 5 × 10^−6^.

Simulated reaches were identified as correct when the trajectory met a sphere with center at [1.7 0 1.3] and radius 0.3. Data points in Figure 3C, D are averages of 15 simulated sessions.

#### LSTM trajectory classification

To classify 3D trajectories as pre- or post-lesion, we employed a Long Short-Term Memory (LSTM) recurrent neural network implemented with python’s PyTorch open-source machine learning library. The input sequence to the network was formed by the time series of 3D coordinates (x, y and z) of each trajectory. We used sequence padding and packing to accommodate variable length trajectories and only process valid non-padded data points. The network architecture comprised the following components: (i) an input layer accepting 3D feature vectors at each time step, (ii) an LSTM module with 2 stacked recurrent layers, each containing 64 hidden units with dropout regularization (0.1) applied between layers to prevent overfitting, which captures temporal dependencies in the sequential input patterns, (iii) a fully connected output layer with sigmoid activation that transforms the final hidden state into a binary classification probability and (iv) binary cross-entropy loss for optimization. No additional activation functions (e.g., ReLU) were required between the LSTM and output layers, as the LSTM units inherently handle non-linearities by incorporating sigmoid and tanh activations. A unidirectional LSTM architecture was chosen as natural movement trajectories are causally produced in time. A bidirectional design would violate the causality of these signals by using future trajectory information to interpret past kinematics. The final classification decision was derived from the hidden state at the last time sample of each sequence, with a decision threshold of 0.5 applied to the output probability.

We pooled trajectories from all mice and sessions. Each session was truncated so that it contributed the same number of trials (N=70) to the data pool. Pooled data were split into training (60%), validation (20%), and test (20%) sets using stratified sampling to maintain class balance across sets. Models were trained using the Adam optimizer with 0.002 learning rate and up to 100 epochs with early stopping based on validation performance. Optimal hyperparameters (number of LSTM layers, number of hidden units, dropout rate, learning rate and optimizer) were identified by performing a grid search across all parameters and selecting the combination yielding the highest validation accuracy.

We trained and evaluated separate models using optimal hyperparameters on the reach, graps, retract, full, full without reach and full without retract trajectory components. Each component was normalized to the [-1, 1] range and processed as an independent dataset. For statistical comparison, we repeated the training and evaluation procedure 199 times for each dataset using different random train, validation and test splits (stratified by class label).

#### Statistics

To compare behavioral metrics between pre- and post-lesion conditions, we employed a Linear Mixed-Effects (LME) model (*fitlme.m*, Matlab). This approach was necessary to account for the hierarchical structure of our dataset, in which multiple measurements (three sessions per condition) were nested within individual mice. Unlike a standard t-test, the LME model accounts for the non-independence of the three repeated measures from each mouse and controls for inter-subject variability by treating the lesion condition (pre vs. post) as a fixed effect and the subject as a random effect (random intercept). Consequently, the model accounts for both inter-mouse variability and the correlation of measurements within individual mice across sessions.

For all other statistical comparisons (Fig. 1) normality was evaluated using the Lilliefors test. We applied parametric Student’s t-tests to normally distributed data and non-parametric Wilcoxon signed-rank tests to datasets that violated normality assumptions.

To identify significant changes in the tuning properties of individual neurons (Fig. 1E,F), we employed a bootstrap-based permutation test. For each neuron, tuning curves were re-computed 1999 times using trials randomly sampled with replacement. We then calculated the distribution of differences between early and late sessions. A change was deemed significant (p<0.05 or p<0.01) if the 95th or 99th percentile confidence intervals of the bootstrapped differences excluded zero.

---

**Figure S1, related to Figure 2.**
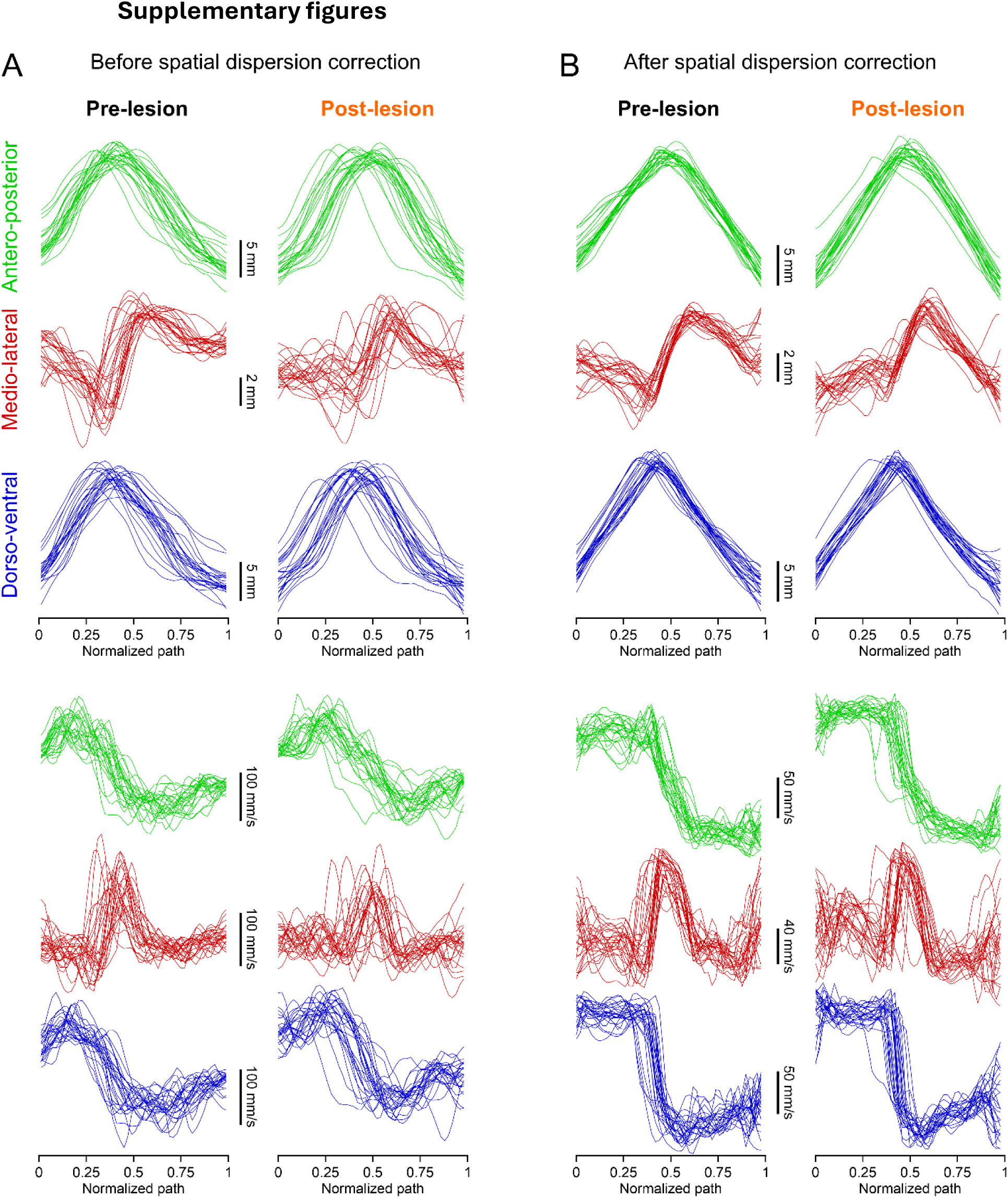
**A**: Position and velocity time series plots of normalized trajectories along the three movement axes of the pre- and post-ablation example sessions in Fig. 2C (only the first 25 trials of the session are shown for clarity). **B**: Same data as in A after spatial trajectory alignment.

**Figure S2, related to Figure 2.**
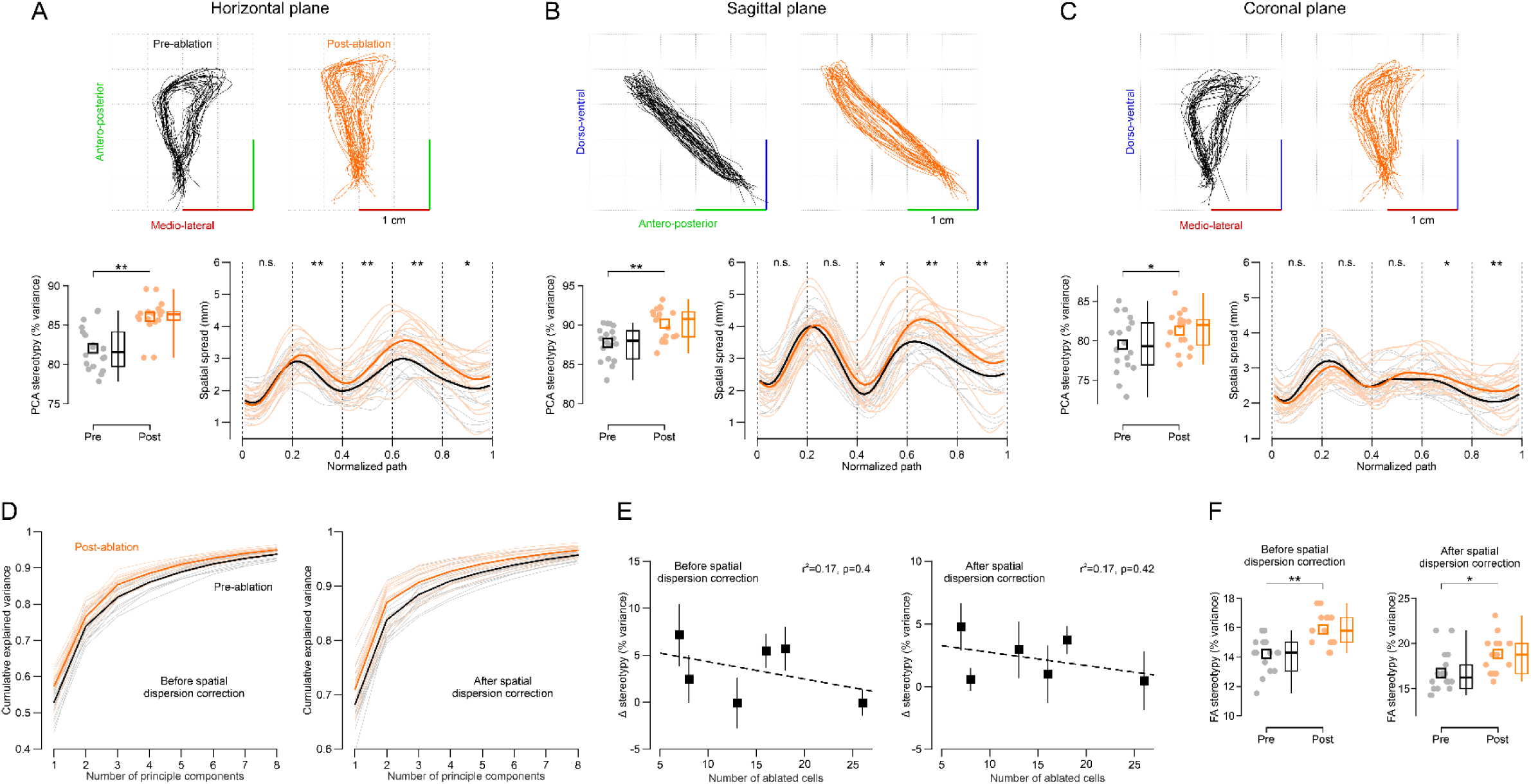
**A-C**: Comparison of pre- and post-ablation trajectory stereotypy and spatial dispersion in the 3 planes of motion (horizontal, sagittal and coronal). Top: same example sessions as in Fig. 2C. **D**: Cumulative explained variance as a function of number of principle components used for the PCA stereotypy analysis for movement trajectories before and after spatial alignment. **E**: n.s.: p>0.05, *: p<0.05, **: p<0.01 (linear mixed effects model fit).

**Figure S3, related to Figure 4.**
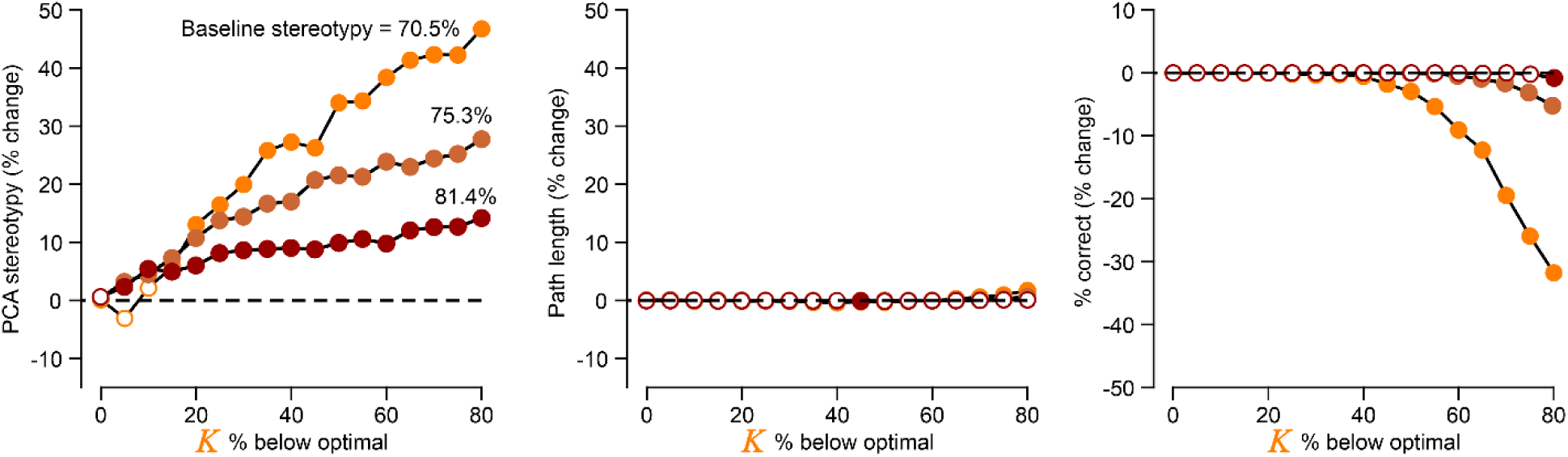
Simulated changes in stereotypy, path length and correct performance with respect to optimized parameters (zero, black dotted line) for different levels of downscaled K, when the model was initialized with three different levels of motor and sensory noise [σ, σ_s_] = [0.1, 0.3], [0.06, 0.2] and [0.04, 0.12] yielding increasing levels of baseline stereotypy.

