## Supplementary figures and images for "Proprioceptive Cortical Neurons Implement Optimal State Estimation"

### Figure S1

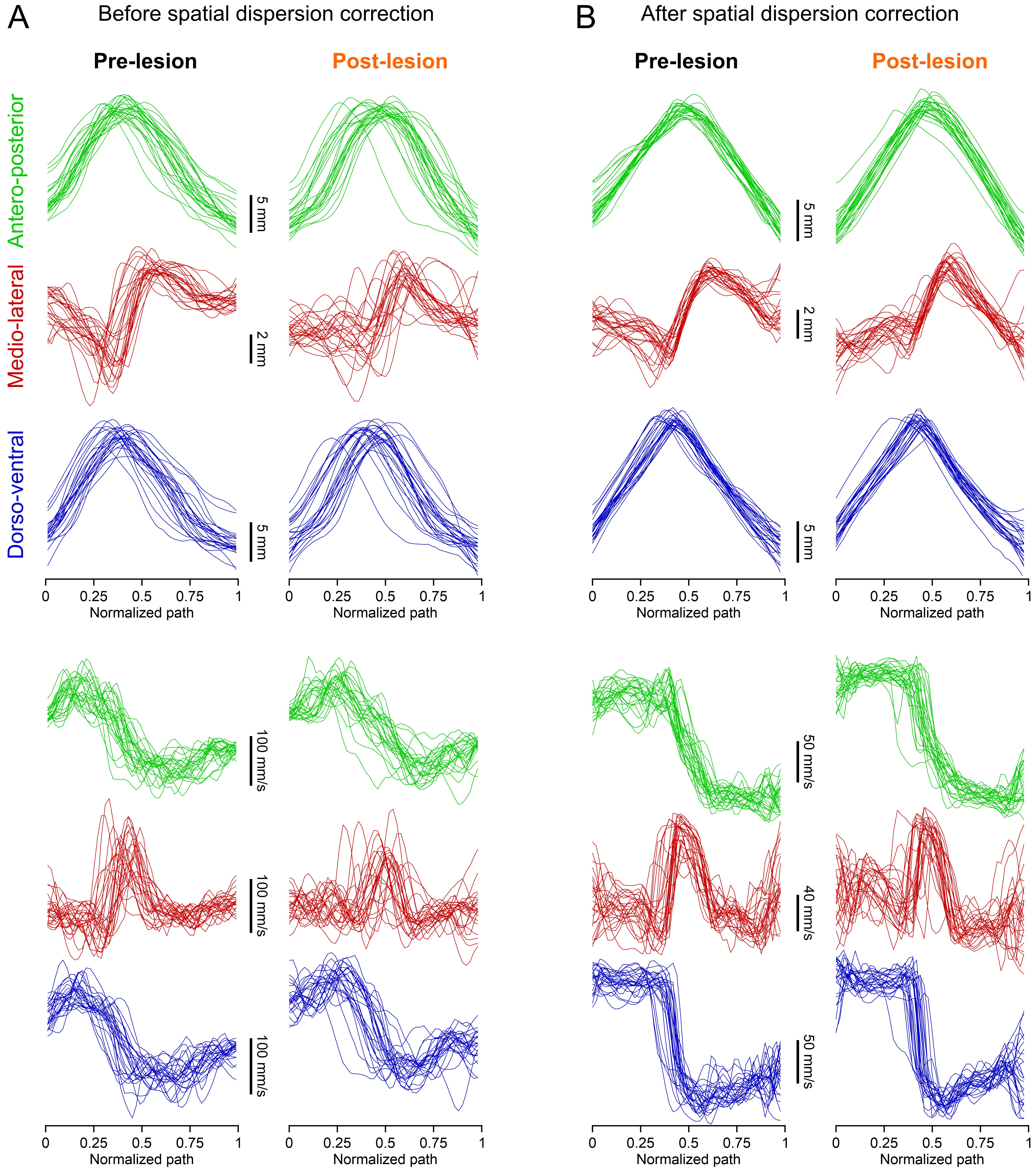

### Figure S2

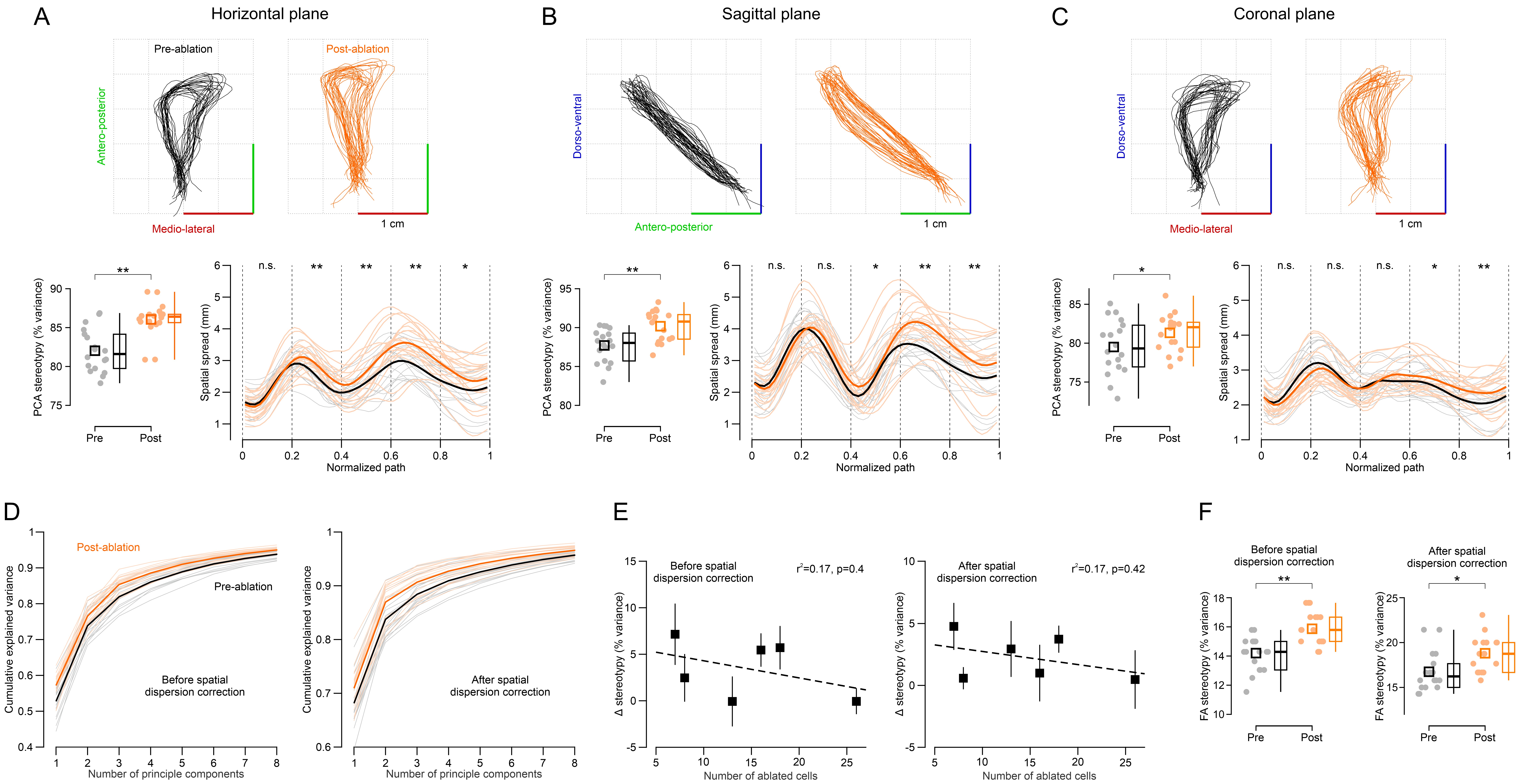

### Figure S3

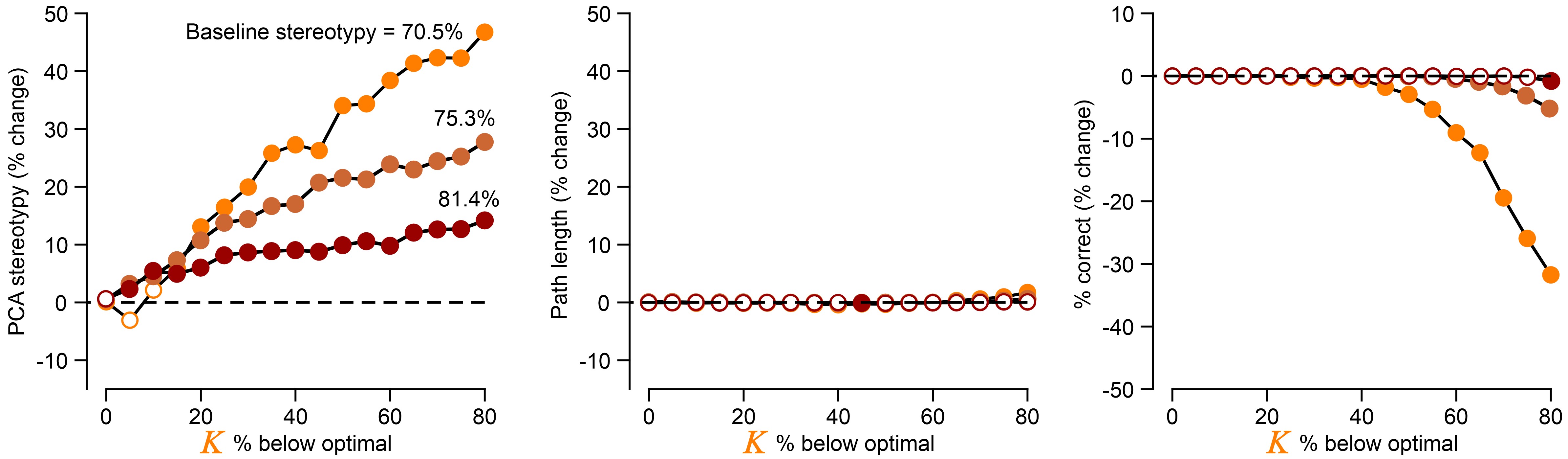
